# Spatial Transcriptomic Analysis of Myositis Muscles Reveals Novel Molecular Insights of MDA5+ Dermatomyositis

**DOI:** 10.64898/2026.01.04.696800

**Authors:** Yulong Qiao, Gexin Liu, Michael Tim-Yun Ong, Hao Sun, Ho So, Huating Wang

## Abstract

Idiopathic inflammatory myopathies (IIM), also known as myositis, are a rare and heterogeneous group of autoimmune disorders characterized by chronic inflammation of skeletal muscle. Other organs are also frequently affected, such as skin, joints, and lungs, leading to morbidity and even mortality. Anti-MDA5 autoantibody-positive dermatomyositis (MDA5+ DM) is a unique subtype with a high mortality rate but the underlying molecular mechanisms remain incompletely understood. In this study, we profiled the spatial transcriptome of a cohort of healthy controls, MDA5+, and MDA5- muscles and found a distinct transcriptomic profile of the MDA5+ muscles with a low percentage of stressed myofibers but increased immune cell infiltration. Niche analysis revealed a distinct microenvironment in the MDA5+ muscles with the endothelial cell (EC) niche showing increased inflammation, dampened oxygen transport function, and a unique enrichment of the Type 1 interferon response. Cell-cell communication and pathway activity analysis uncovered that hypoxia, TGF-β, NF-κB, and TNF-α signaling were enriched in the MDA5+ EC niche. Altogether, our results support a vasculopathy model whereby blood vessels exhibit a strong inflammatory response and impaired oxygen transport function, leading to vasculopathy and perivascular immune cell infiltration in MDA5+ muscles.

## Introduction

Idiopathic inflammatory myopathies (IIM), also known as myositis, are a rare and heterogeneous group of autoimmune disorders characterized by immune-mediated muscle damage. Besides muscles, other organs are frequently affected, such as skin, joints, lungs, and gastrointestinal tract, leading to morbidity and even mortality. Based on clinical, histopathological, and serological features, myositis can be classified into several subgroups: dermatomyositis (DM), clinical amyopathic dermatomyositis (CADM), polymyositis (PM), overlap myositis (OM), immune-mediated necrotizing myopathy (IMNM), anti-synthetase syndrome (AsyS), and inclusion body myositis (IBM)^1^. The discovery of myositis-specific autoantibodies (MSAs), which are exclusively found in patients with myositis and correlate with specific clinical phenotypes, improved the diagnosis and classification of patients into more homogeneous groups, outcome prediction, treatment strategies, as well as understanding of molecular mechanisms^2^. For example, anti-TIF1 (Transcriptional intermediary factor 1) and anti-Mi-2 autoantibodies are specifically associated with DM. Patients with anti-TIF1 autoantibodies often present with mild muscular symptoms, but are likely to have dysphagia; adult patients with anti-TIF1 autoantibodies have a strong possibility of developing malignancies. In contrast, patients with anti-Mi2 antibody-positive DM often experience severe muscle involvement, characterized by muscle weakness, elevated creatine kinase (CK) levels, and myofiber degeneration/necrosis that appears at perifascicular sites on muscle biopsies.

Anti-MDA5 (melanoma differentiation-associated gene 5) autoantibody-positive DM constitutes a distinct subtype of myositis. It is characterized by skin ulceration and rapidly progressive interstitial lung disease (RP-ILD). RP-ILD, defined as progressive dyspnea, progressive hypoxemia, and worsening of interstitial changes on chest radiography within 1 month from the onset of respiratory symptoms, is the most severe manifestation, associated with a high mortality rate of approximately 50%^2^. Interestingly, there is minimal or absent muscle involvement in the MDA5+ patients. Their muscle biopsies appear normal or show only minimal anomalies, with scarce inflammation, focal perimysial macrophage infiltration, absent or focal MHC-I expression, and rarely observed complement deposition, compared to other DM serotypes^3, 4^. Of note, the prevalence of MDA5+ myositis is significantly higher in East Asian populations compared to European cohorts, where it is markedly associated with interstitial lung disease (ILD) and a high frequency of RP-ILD. A cohort from Hong Kong, for instance, found that 30% of DM patients were MDA5+, and two-thirds of MDA5+ patients had RP-ILD^2,5^. The current treatment of MDA5+ myositis with conventional glucocorticoids and immunosuppressants shows limited efficacy, resulting in a poor overall prognosis. This highlights the critical need to investigate the underlying molecular underpinnings of MDA5+ myositis and to explore innovative therapeutic strategies for patients.

The precise aetiology and pathogenesis of MDA5+ myositis remain incompletely understood. The disease is thought to be initiated by complex interactions between genetic susceptibility and environmental triggers, which break immune tolerance and lead to the production of anti-MDA5 autoantibodies. Multiple factors, including anti-MDA5 autoantibodies, immune cells, and pro-inflammatory mediators, are implicated in the immunopathogenesis of MDA5+ myositis^6^. A hallmark of this condition is a pronounced upregulation of type I interferon (IFN-I)-inducible genes—the so-called “IFN-I signature”—observed in the skin, lungs, muscle, and peripheral blood. This signature correlates with the progression of ILD and poorer prognosis, implicating the role of IFN-I in pulmonary pathology^7^. Furthermore, severe vasculopathy is a characteristic histopathological finding in skin biopsies from MDA5+ patients. Evidence suggests IFN-I drives this endothelial injury, as demonstrated by elevated expression of STAT1 and the interferon-inducible genes MX1 and ISG15 within cutaneous blood vessels^8, 9^. In skeletal muscles, the IFN-I signature is upregulated in DM, including MDA5+ patients. However, MDA5+ patients often exhibit mild or absent clinical muscle weakness, and biopsy results show no or minimal atrophy. It remains unclear why the skeletal muscle is relatively spared from severe damage, whereas the lungs and skin are frequently affected in MDA5+ myositis.

Recent advances in single-cell(sc) level profiling have enabled in-depth cellular/molecular characterization of myositis. scRNA-seq studies of peripheral blood mononuclear cells (PBMCs) from MDA5+ patients revealed increased proportions of CD14+ monocytes, plasma cells, peripheral antibody-secreting cells, and CD8+ T cells, alongside an overactivated IFN-I response in these immune cells^10, 11^. However, how the immune dysfunction and immune cells mediate the inflammation and affect the muscles of MDA5+ patients remains largely unknown. Spatial transcriptomics enables the analysis of gene expression within its tissue context that is lost in single-cell sequencing. Several recent studies applying spatial transcriptomics on IBM and juvenile dermatomyositis (JDM) muscle biopsies discovered the selective type 2 fiber vulnerability linked to genomic stress and denervation pathways in IBM^12^, as well as mitochondrial abnormalities and an upregulated IFN-I signature in the atrophy region of JDM^13,14^.

In this study, we performed spatial transcriptomic profiling on muscle biopsies from 3 healthy controls and 11 myositis patients, including 3 MDA5+ and 8 MDA5-. Our findings reveal that despite the absence of clinical muscle weakness, MDA5+ muscles exhibited a distinct spatial architecture characterized by a robust IFN-I signature and tissue microenvironment remodeling. While exhibiting a lower burden of stressed myonuclei, MDA5+ muscles showed evident immune infiltration in the perivascular region. Six distinct niche types were identified within myositis muscles, and the EC niche in MDA5+ muscles uniquely demonstrated an upregulation of inflammatory and IFN-I response alongside a downregulation of oxygen transport pathways. Cell-cell communication analysis further revealed heightened hypoxia, TGF-β, TNF-α, and NF-κB signaling, which were particularly enriched within the MDA5+ EC niche. Moreover, functional analysis of immune cells implicated macrophages in driving localized humoral immune responses. Collectively, these findings for the first time delineate a unique spatial immunopathology in MDA5+ muscles and reveal potential therapeutic targets.

## Results

### Spatial transcriptomics reveals a distinct gene expression profile in MDA5+ muscles

To uncover molecular insights underlying the pathogenesis of MDA5+ myositis in skeletal muscle, we performed spatial transcriptomic sequencing of muscle biopsies collected from myositis patients at the Prince of Wales Hospital, Hong Kong (Fig. 1A). Eleven patients were included in the study, including 4 cases of CADM (3 MDA5+), 3 OM, 2 DM, 1 PM, and 1 IMNM. Hamstring muscles collected from patients undergoing knee replacement surgery were used as control (Ctrl)^15^ (Suppl Fig. S1A, Suppl Table S1). According to serotypes, the myositis samples were classified as 3 MDA5+ and 8 MDA5- samples (including 2 ro52+, 1 TIF1+, 1 Mi2+, 1 HMGCR+, and 2 seronegative samples). Cryosections of muscles (∼10 µm thickness) were prepared, stained with hematoxylin and eosin (H&E) for imaging, then destained and incubated with Visium human transcriptome probe set. The probes were then transferred to the Visium spatial slides using 10x CytAssist, released, and amplified for sequencing. The data were analyzed using the Space Ranger pipeline, and spots were identified. Spots that failed to cover any tissue area or had fewer than 1000 Unique Molecular Identifier (UMI) counts, fewer than 200 genes, or more than 20% mitochondrial genes were excluded. As a result, a total of 9,112 spots were detected from all samples. The data were of high quality, with 15614 median UMI counts and 3774 median genes per spot detected on average (Suppl Fig. S1B, Suppl Table S1). Spatial Uniform Manifold Approximation and Projection (UMAP) analysis of all samples revealed two distinct clusters (Fig. 1B). Cluster 1 contained the 3 healthy Ctrl samples, while cluster 2 contained the 3 MDA5+ samples (CADM42, CADM63, CADM76), demonstrating a clear separation of the MDA5+ from Ctrl samples (Fig. 1C). The OM (OM49, OM60, OM64), and DM (DM70, DM81) samples, however distributed across both clusters (Fig. 1D), suggesting substantial heterogeneity. Pseudobulk analysis was performed to identify the differentially expressed genes (DEGs) between MDA5+ vs. Ctrl samples. Despite the undetectable muscle weakness, we found immune-related genes such as *ISG15*, *IFI6*, *MX1*, *CMPK2*, *OAS1*, *RSAD2*, etc., were highly enriched in the MDA5+ muscles (Suppl Table S1). The gene ontology (GO) enrichment analysis of the DEGs upregulated in MDA5+ samples also revealed enriched GO terms related to immune responses, such as “defense response to virus”, “negative regulation of viral process”, “cellular response to IFN-I”, etc. (Fig. 1E, Suppl Table S1). The upregulated DEGs in all myositis samples were also enriched for GO terms related to immune responses when compared to the Ctrl samples (Fig. 1F, Suppl Table S1). Similarly, when comparing each subtype to the Ctrl samples, all subtypes were enriched for immune-related pathways (Suppl Fig. S1C-E, Suppl Table S1).

**Figure 1.**
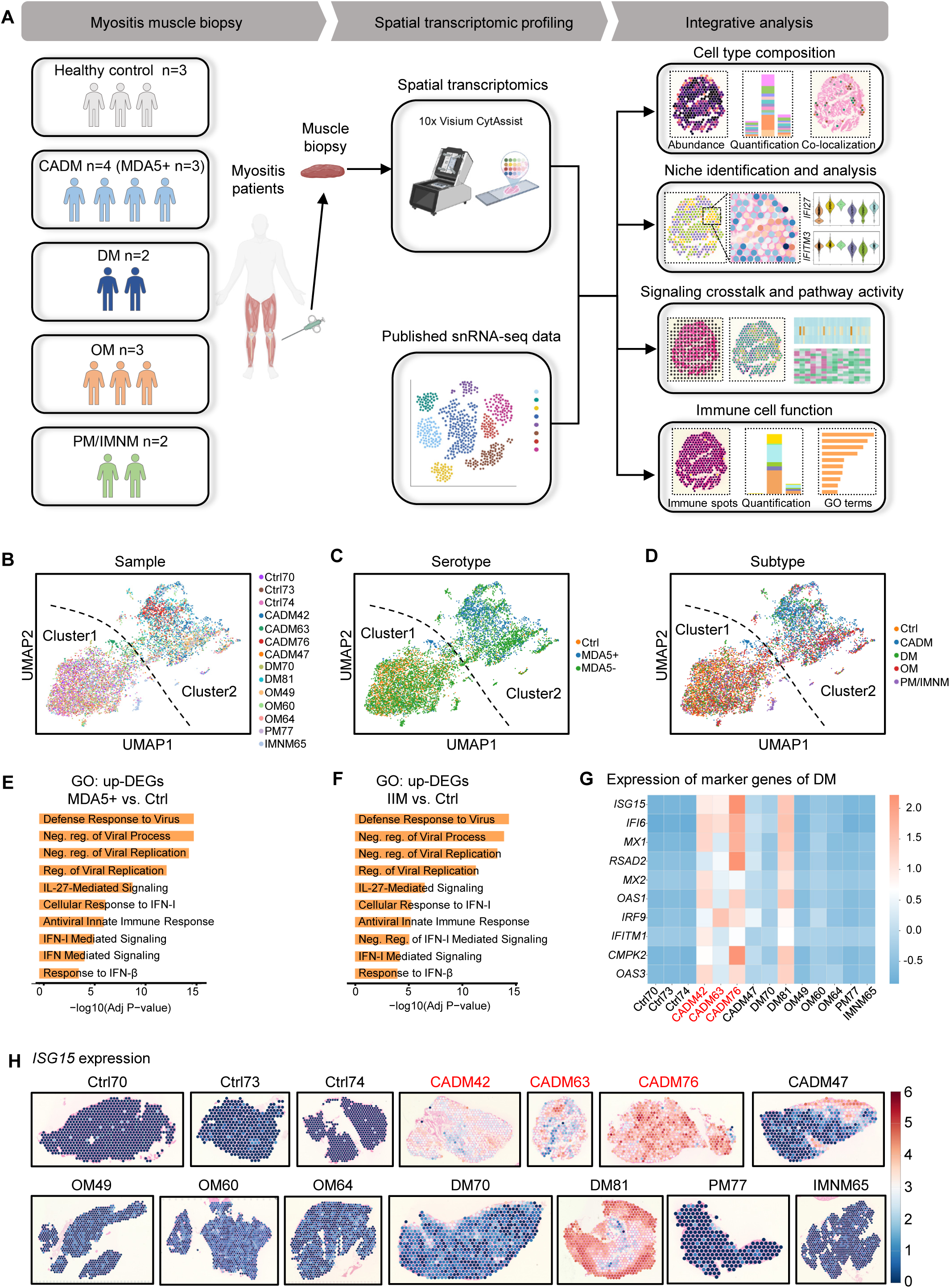
Spatial transcriptomics reveals a distinct gene expression profile in MDA5+ muscles. (A) Schematic showing the overall design of the study. Muscle biopsies were collected from 11 donors with myositis, including 4 CADM (3 MDA5+), 2 DM, 3 OM, and 1 IMNM/1 PM. 3 muscle biopsies from healthy donors were also collected. Spatial transcriptomics profiling was performed using Visium for fresh frozen and CytAssist (10x Genomics). One publicly available snRNA-seq dataset was adopted for cell type deconvolution analysis. Integrated data analysis was performed, including niche identification, niche-enriched DEG analysis, signaling crosstalk, pathway activity, and immune cell spot analysis. (B) UMAP visualization showing Visium spots of each sample. (C) UMAP visualization showing Visium spots of each serotype. (D) UMAP visualization showing Visium spots of each subtype. (E) GO plot of biological process (BP) terms enriched in myositis groups (IIM) using the top 200 upregulated genes, compared to the Ctrl group. (F) GO plot of biological process terms enriched in MDA5+ groups using the top 200 upregulated genes, compared to the Ctrl group. (G) Heatmap showing the mean expression level of DM marker genes in each sample. The mean expression level was transformed into a row z-score for visualization. H) Spatial gene expression of *ISG15* in all samples, raw counts were log-transformed and normalized.

MDA5+ and DM muscles are known to display an IFN-I signature^6^. Indeed, the top 10 frequently upregulated IFN-related genes in DM^16^(*ISG15*, *IFI6*, *MX1*, *RASD2*, *MX2*, *OAS1*, *IRF9*, *IFITM1*, *CMPK2*, *OAS3*) were significantly upregulated in the 3 MDA5+ (highlighted in red) and the DM81 samples(Fig. 1G). For example, a robust *ISG15* upregulation was detected (Fig. 1H). In summary, the spatial transcriptomic analysis of myositis muscles reinforced an increased IFN-I signature in MDA5+ muscles.

### Cell type analysis reveals increased immune cell infiltration in MDA5+ muscles

To further elucidate the pathological alterations in MDA5+ muscles, we dissected the cell type composition of the spatial spots by conducting deconvolution analysis using cell2location^17^; a publicly available single-nuclei RNA-seq data set from myositis muscles was used for cell type annotation^12^. As a result, a total of 22 cell types were identified (Suppl Fig. S1F-H), including major cell types such as Type 1 and Type 2 myonuclei (Type 1/2 MN), stressed myonuclei (sMN), reactive myonuclei (rMN), endothelial cells (ECs), pericytes, fibro-adipogenic progenitors (FAPs), skeletal muscle stem cells (MuSCs), and immune cells. Estimated cell type abundance was quantified, and a high heterogeneity in cell composition was observed among the myositis samples (Fig. 2A, Suppl Table S2). For example, the abundance of sMN (from 20% to 56%) and immune cells (from 1.3%-13%) varied across myositis muscles and within a subtype (20% sMN in CADM63 vs. 45% in CADM76, and 32% sMN in OM49 vs. 56% in OM64) (Fig. 2A). When comparing the 3 MDA5+ with 8 MDA5- muscles we found a lower percentage of sMNs in the MDA5+ DM group (30% vs. 41%), which is in line with the undetectable muscle weakness and low CK level in the MDA5+ patients (Suppl Table S1). Surprisingly, an increased percentage of immune cells was found in the MDA5+ vs MDA5-muscles (10% vs 4%) (Fig. 2B). For instance, as shown in Fig. 2C, the MDA5+ CADM76 muscle showed a reduced abundance of sMN and an increased abundance of B cells, plasma cells, and CTLs compared to the MDA5- muscles (DM70 and OM64). Among different subtypes (4 CADM, 3 OM, 3 DM, and 2 IMNM/PM), OM and DM muscles showed a relatively higher abundance of sMN (Fig. 2D).

**Figure 2.**
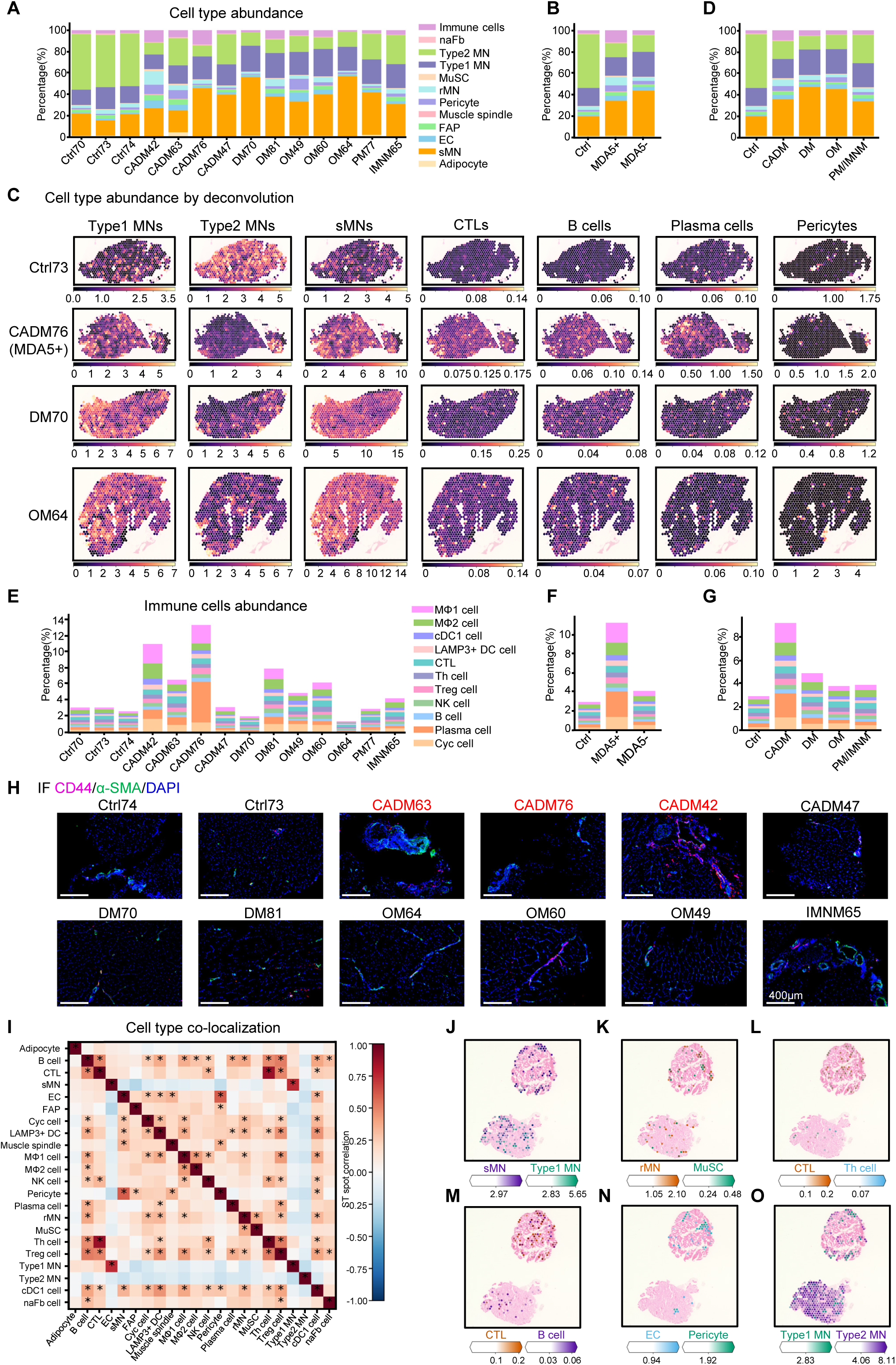
Cell type analysis reveals increased immune cell infiltration in MDA5+ muscles. (A) Cell type composition in each sample, calculated by deconvolution of cell types. (B) Comparison of cell type composition in Ctrl, MDA5+, and MDA5- groups. (C) Cell type composition of Type1 myofiber nuclei (Type1 MN), Type 2 myofiber nuclei (Type2 MN), stressed myofiber nuclei(sMN), Cytotoxic T cells (CTLs), B cells, plasma cells, and pericytes in 4 representative samples. (D) Comparison of cell type composition in myositis subtypes. (E) Abundance of immune cell types in each sample. (F) Comparison of immune cell abundance in Ctrl, MDA5+, and MDA5- groups. (G) Comparison of immune cell abundance in each myositis subtype. (H) Immunofluorescence staining of CD44(red) in representative samples. Nuclei were stained with DAPI (blue), and blood vessels were stained with α-SMA (green). Scalebar = 400 μm. (I). Heatmaps showing the correlation of spatial locations of cell types (*, Pearson correlation coefficient > 0.25). (J-N) Spatial plots showing the abundance and co-localization of sMNs&Type1(J), rMN&MuSC(K), CTL&Th cell(L), CTL&B cell(M), EC&pericyte(N) in Ctrl73 and CADM63 samples. (O) Spatial plots showing the abundance and separate distribution of Type 1 MNs and Type 2 MNs in Ctrl73 and CADM63 samples.

We then examined the immune cell types closely, a total of 11 types of immune cells were identified including type 1 and type 2 macrophages (MΦ1, MΦ2), canonical type 1 dendritic cells (cDC1 cells), LAMP3+ dendritic cells (LAMP3+ DC cells), cytotoxic T cells (CTLs), helper T cells (Th cells), regulatory T cells (Treg cells), natural killer cells (NK cells), B cells, plasma cells, and cycling immune cells (Cyc cells) (Fig. 2E). An increase in the immune cells was observed in the MDA5+ vs MDA5- group (Fig. 2F, Suppl Fig. S2A). For instance, the percentage of MΦ1, MΦ2, plasma cells, and Cyc cells showed 3.5-, 2.2-,6.4-, and 2.5-fold increase, respectively (Suppl Fig. S2A). The CADM subtype also showed an increase in MΦ1, plasma cells, and Cyc cells, compared to DM and OM muscles (Fig. 2G, Suppl Fig. S2B). To validate the above finding, we stained the CD44 (marker for immune cells and infiltration) and found increased signals in the myositis muscles, especially in the vascular regions of the two MDA5+(CADM63, CADM42) and OM60 muscles (Fig. 2H).

Furthermore, we dissected the spatial co-localization of cell types across all samples (Fig. 2I) and found that sMN and Type I MN frequently co-occupied the same spatial spots, suggesting that Type I fibers may be particularly sensitive to stress (Fig. 2I, 2J). MuSCs frequently co-localized with rMN, highlighting their role in repairing the muscle fibers (Fig. 2I, 2K); Th cells were found to co-localize with CTLs, possibly facilitating CTL activation.^18^ (Fig. 2I, 2L); B cells also exhibited a significant co-localization with CTLs (Fig. 2I, 2M) alongside Th and cDC1 cells, indicating a complex immune niche where these cells engage in crosstalk, maturation, and activation^18, 19^; Expectedly, ECs and pericytes co-localized at the blood vessels (Fig. 2N). Type 1 and Type 2 MN displayed a negative correlation and were distributed separately (Fig. 2O). In summary, the above results reveal an increase in immune cells, accompanied by a low abundance of sMN in MDA5+ muscles.

### Niche analysis reveals a distinct cellular microenvironment in MDA5+ muscles

To further dissect the complex microenvironment in the myositis muscles, we then identified cellular niches defined by the cell type composition of spatial spots. Non-negative matrix factorization (NMF) analysis was conducted based on the deconvoluted cell types of each spot (Fig. 3A). As a result, 6 types of niches were identified, the “EC niche” (dominated by ECs and pericytes), “FAP niche” (dominated by FAPs), “rMN niche” (consisted of rMN, MuSCs and immune cells), “Type 1 MN niche” (dominated by Type 1 MN), “Type 2 MN niche” (dominated by Type 2 MN), and “sMN niche” (consisted of sMN and immune cells such as B cells, plasma cells, and Treg cells) (Fig. 3B, C). For instance, the quantification of cell compositions of sMN niches showed dominance by sMNs (Fig. 3D-E). Furthermore, DEG analysis revealed distinct expression profiles of each niche type, and the top-ranked DEGs were consistent with their cell type compositions (Fig. 3F, Suppl Table S3). For example, EC markers *VWF*, *CDH5*, and *PECAM1*, smooth muscle cell markers *MYH11*, *ACTA2*, and *TAGLN* were highly expressed in the EC niche; FAP markers *DCN*, *FBN1*, *MFAP5*, and *PCOLCE* were highly expressed in the FAP niche; Muscle regeneration related genes *MYOG* and *MYH3* were highly expressed in the rMN niche; Autophagy related gene *SQSTM1*, stress response genes *MKNK2*, *MTNL*, *SELENOM*, *CRYAB*, and *NFE2L1* were upregulated in the sMN niche; Fast skeletal muscle fiber associated genes *MYH1*, *MYH2*, *TNNT3*, and *TNNI2* were enriched in the Type 2 MN niche while slow-twitch fiber associated genes *MYH7B*, *MYH7*, *ATP2A2* were enriched in the Type 1 MN niche (Fig. 3F). Expectedly, a limited number of sMN niches were detected in the 3 Ctrl muscles, but a much higher number were found in the myositis muscles (Fig. 3G, H). A lower percentage of sMN niches was also detected in the MDA5+ vs MDA5- muscles (33% vs. 41%) (Fig. 3I). When comparing subtypes, a significantly higher proportion of the sMN niche was observed in DM (58%) and OM (50%), compared to 31% in CADM (Fig. 3J). The niche heterogeneity was also detected within subtypes. For instance, 25% of the sMN niches were found in CADM47 vs. 4% in CADM63, and 31% in DM81 vs. 86% in DM70 (Fig. 3G, H). In summary, we identified 6 distinct cellular niches in myositis muscles, and MDA5+ muscles displayed a unique microenvironment composed of a low proportion of the sMN niches.

**Figure 3.**
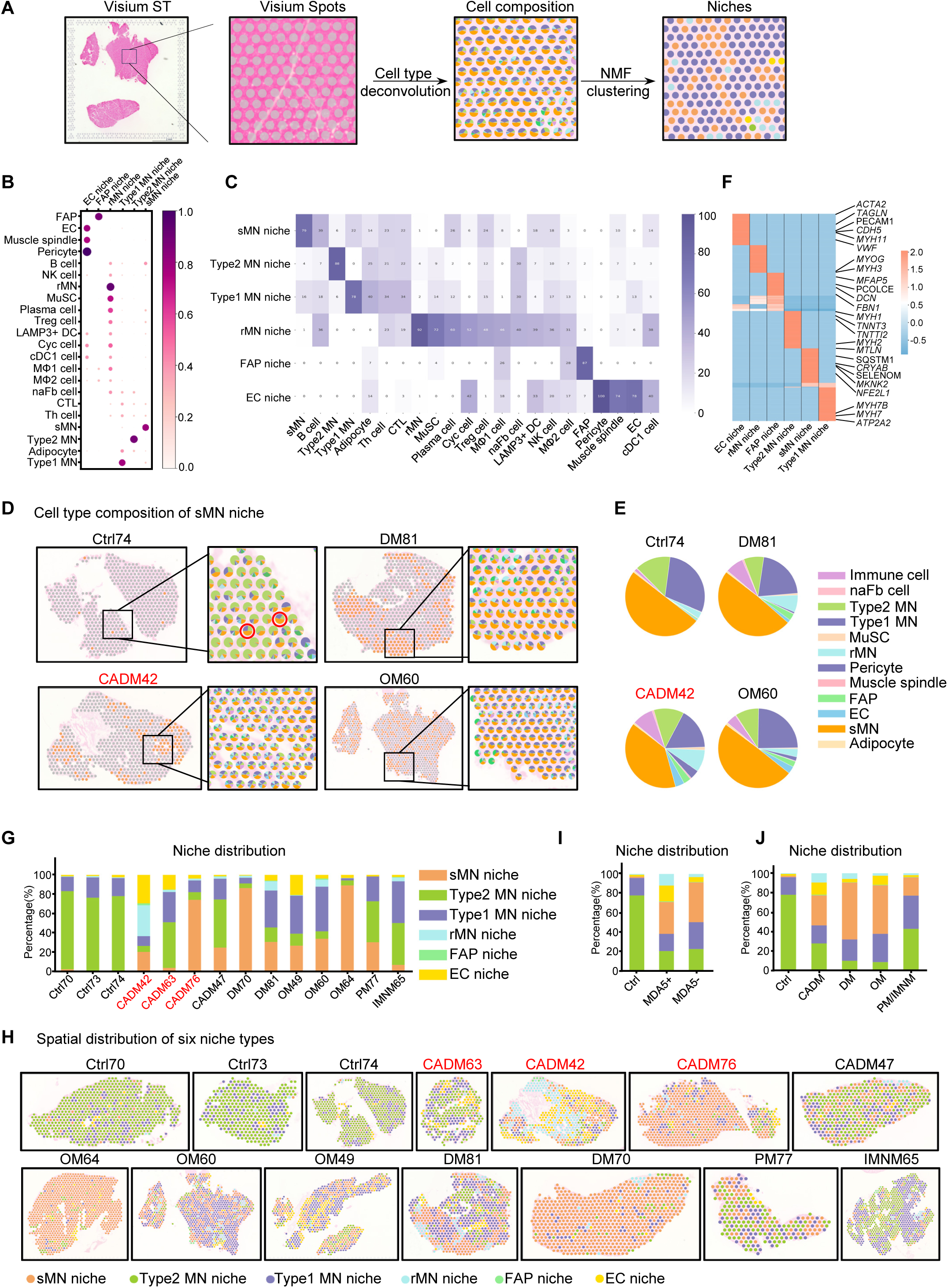
**Niche analysis reveals a distinct cellular microenvironment in MDA5+ muscles. (**A) Schematic illustration of the niche identification with non-negative matrix factorization (NMF) analysis based on cell type composition. (B) Dot plot of the estimated NMF weights of cell types (rows) across niche types (columns). (C) Heatmap showing the normalized distribution of estimated cell type abundance in percentages across the niches. (D) Illustration of the “sMN niche” dominated by sMNs in four representative samples. (E) Pie chart showing the cell type composition of the “sMN niche”. (F) Heatmap showing the top-ranked 50 genes in each niche type. (G) Bar plot shows the niche composition in all samples. (H) Spatial plot showing the distribution of six types of niches across all samples. (I) Bar plot comparison of the niche composition in Ctrl, MDA5+, and MDA5- groups. (J) Bar plot comparison of the niche composition in each subtype.

### Unique gene expression in the EC niche of MDA5+ muscles

To further dissect the molecular changes of the MDA5+ muscles, we analyzed DEGs in the above-defined niches (Suppl Table S4). When comparing type 1 and type 2 MN niches in the MDA5+ vs. MDA5- muscles, 332 and 1725 DEGs were identified, respectively; the up-DEGs were both enriched for GO terms related to “cellular response to IFN-I”, “IFN-I mediated signaling pathway”, “defense response to virus”, “IL 27-mediated signaling pathway”, “antiviral innate immune response”, etc., and the down-DEGS were enriched for “neuromuscular junction development”, “actomyosin organization”, etc. (Fig. 4A, B). 735 DEGs were identified in the MDA5+ vs. MDA5- sMN niches, and the up-DEGs were also enriched for similar IFN-related GO terms (Fig. 4A). For example, IFN-stimulated gene, *OASL*, was upregulated in type 1 MN, type 2 MN, and sMN niches in the MDA5+ muscles (Fig. 4C). *NUPR1*, a stress-response gene that is a critical repressor of ferroptosis^20, 21^, was upregulated in all myositis samples but showed a higher expression in the MDA5+ muscles (Suppl Fig. S3A). When comparing the rMN niches, 655 DEGs were identified in the MDA5+ vs MDA5- muscles, and the up-DEGs were dominated by “extracellular structure organization”, “extracellular matrix organization”, “collagen fibril organization”, etc. (Suppl Fig. S3B). The up-DEGs in the FAP niches were enriched for “ECM pathways” while “sarcomere organization” and “myofibril assembly” related pathways were downregulated (Suppl Fig. S3C).

**Figure 4.**
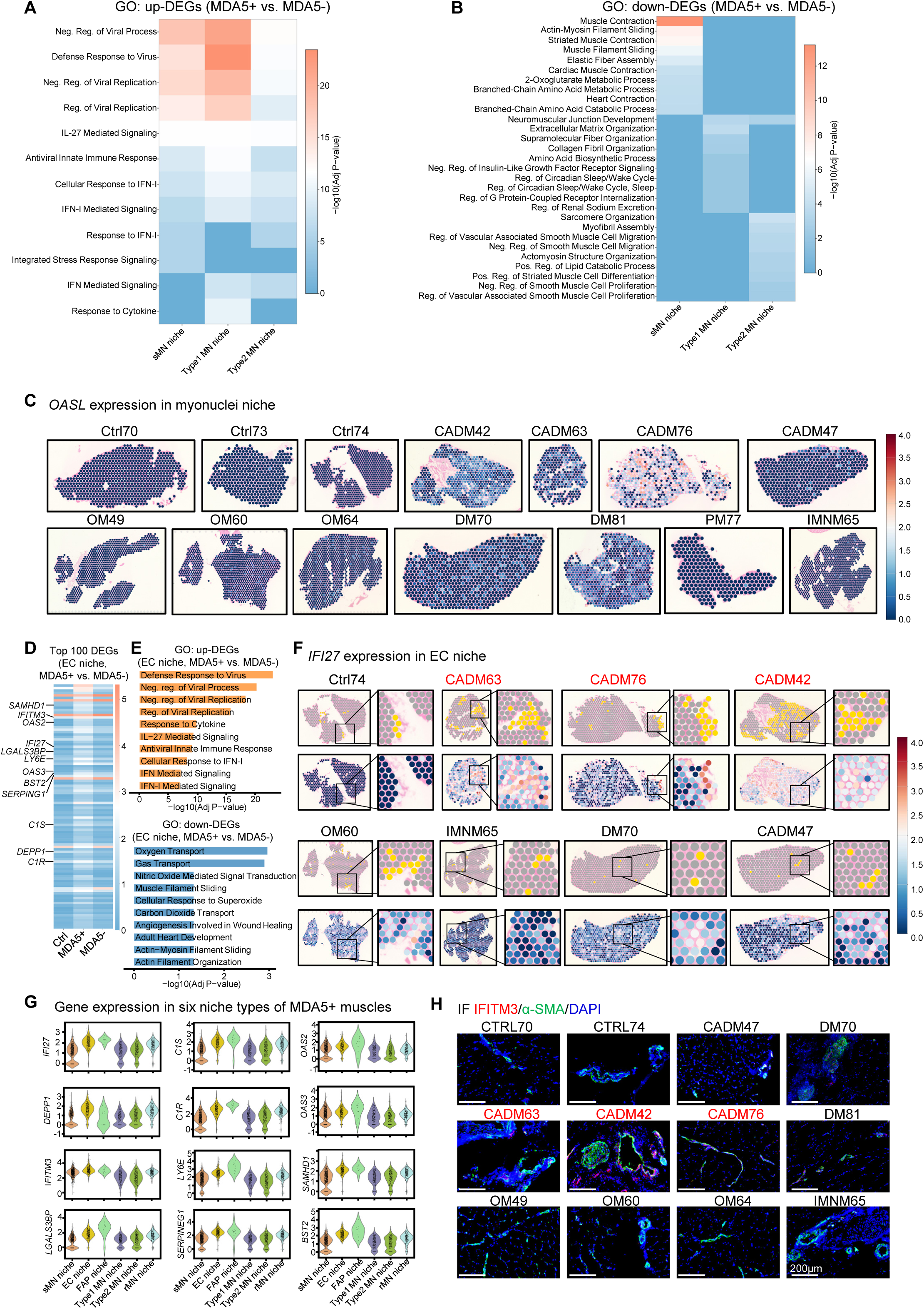
**Unique gene expression in the EC niche of MDA5+ muscles. (**A) Heatmap showing the top 10 enriched GO terms of upregulated genes in the sMN niche, type1/2 MN niche, colored by–log10 P value. (B) Heatmap showing the top 10 enriched GO terms of down-regulated genes in the sMN niche, type1/2 MN niche, colored by–log10 P value. (C) Normalized spatial expression of *OSAL* in all samples. (D) Heatmap showing the top 100 DEGs in the EC niche of Ctrl, MDA5+, and MDA5- groups. (E) Top 10 enriched GO terms of up- and down-regulated genes in the EC niche of MDA5+ vs MDA5- muscles. (F) Spatial plots showing the distribution of EC niches (yellow spots) and the expression of *IFI27* in representative Ctrl, MDA5+, and MDA5- muscles. (G) Violin plots showing the expression of genes enriched in the EC niche of MDA5+ muscles. (H) Immunofluorescence staining of IFITM3 (red) and α-SMA (green) in representative myositis muscles. Nuclei were stained with DAPI (blue). Scalebar = 200 μm.

When comparing the MDA5+ vs. MDA5- EC niches, 613 DEGs were identified, and the up-DEGs were enriched for anti-viral responses and responses to IFN-I related pathways, while the “oxygen transport”, “gas transport”, “Nitric Oxide Mediated Signal Transduction”, and “carbon dioxide transport” pathways were enriched in the down-DEGs, indicating increased immune response and dampened oxygen/gas exchange function of blood vessels of the MDA5+ muscles (Fig. 4D, E). For example, *IFI27* gene expression was upregulated in MDA5+ vs. MDA5- EC niches (Fig. 4D, F). Of note, the expression of *IFI27* also showed an enrichment in the EC niches compared to other myonuclei niche types in the MDA5+ muscles (Fig. 4F, G). Similarly, a group of up-DEGs in the MDA5+ EC niche, including IFN-I-stimulated genes (*IFITM3*, *OAS2*, *OAS3*, *LY6E*, *BST2*), anti-viral immune-related genes (*LGALS3BP* and *SAMHD1*), complement system-related genes (*SERPING1*, *C1S*, *C1R*), and autophagy-related gene *DEPP1*, were also enriched in the EC niche of MDA5+ muscles, compared to other myonuclei niche types of MDA5+ muscles (Fig. 4G). This was further validated by immunofluorescence (IF) staining of IFITM3, SAMHD1, IFI27, and BST2, co-staining with α-SMA to label the blood vessels; increased staining was found in the blood vessel regions in the MDA5+ muscles (Fig. 4H, Suppl Fig.S3D-F). We then examined the source of IFNs that activate the expression of these IFN-I inducible genes in MDA5+ muscles. Surprisingly, Type I, II, and III IFN transcripts were barely detectable in the Ctrl or myositis muscles, but several IFN receptors, such as *IFNAR1/2* and *IFNGR1/2*, were upregulated (Suppl Fig. S4G).

The transcriptomic difference of each niche type was also dissected among OM and DM subtypes. The OM muscles showed a reduced response to IFN-I in the sMN, EC, type 1, and rMN niches, compared to other subtypes. While the DM muscles exhibited the known IFN-I signature in the EC, type 1, and type 2 niches (Suppl Fig. S4). Taken together, the analyses discovered a cohort of proinflammatory and IFN-I inducible genes enriched in the EC niche of MDA5+ muscles, highlighting the prominent IFN-I signature and vasculopathy of MDA5+ muscles.

### Cellular crosstalk analysis uncovers enriched signaling pathway activities in the EC niche of MDA5+ muscles

Next, to dissect the cell-cell communication between niches, we analyzed the crosstalk signals using the COMMOT method^22^. 42 signaling pathways were detected, and most of them were heightened in myositis muscles (Fig. 5A, Suppl Fig. S5A, Suppl Table S5). Among these signaling pathways, ACTIVIN^23–25^, ANGPTL2^26^, and TWEAK are related to muscle atrophy; CCL^27^, CSF^27^, galectin-9-cd44^28, 29^, CHEMERIN^30^, MIF, and WNT ^31^ promote inflammation; COMPLEMENT^32, 33^ is related to innate immune responses; cytokines CX3C^34^, CXCL^27, 29^, GDF11^35^ are known to promote muscle atrophy; granulin(GRN), is reported to be upregulated in DM serum^36^; Midkine (MK) signaling is upregulated in muscle regeneration^37–39^ and can also promote recruitment of immune cells and be involved in autoimmune diseases^40^; PDGF, is related to muscle regeneration and inflammation^41^; PERIOSTIN signaling is related to fibrosis and inflammation^42^; VEGF signaling and may promote the pathogenesis through forming blood vessels and increasing the immune cell infiltration^41, 43, 44^; Visfatin signaling is upregulated in myositis serum and muscles, which may promote inflammation^45, 46^. Of note, TGF-β signaling was evidently upregulated in 7 of 11 myositis muscles (Fig. 5B) and higher in the MDA5+ vs. MDA5- muscles (Fig. 5C). Furthermore, we found sMN niches secreted the highest amount of TGF-β signaling molecules (Fig. 5D), consistent with the report that TGF-β signaling could be secreted from “inflammatory myonuclei”^47^. We also analyzed the published scRNA-seq data^12^ using CellChat, which consistently revealed extensive cellular crosstalk mediated by TGF-β, MIF, COMPLEMENT signaling, etc. (Suppl Fig. S6).

**Figure 5.**
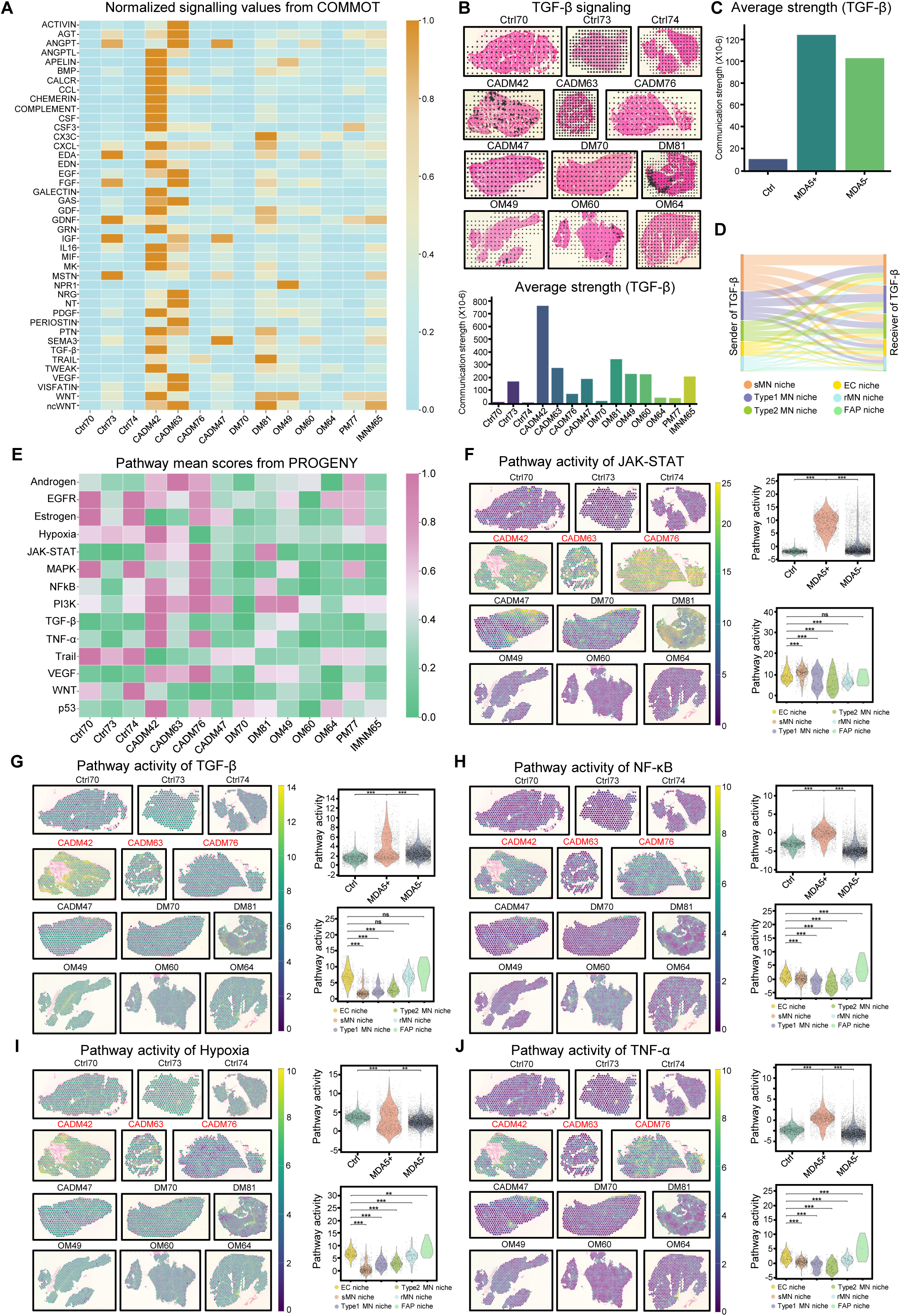
**Cellular crosstalk analysis uncovers enriched signaling pathway activities in the EC niche of MDA5+ muscles. (**A) Heatmap showing the normalized signaling communication strength calculated by COMMOT in all samples. (B) The signaling direction (top) and the normalized signaling communication strength (bottom) in each sample are shown for TGF-β signaling. (C) Comparison of the normalized signaling communication strength for TGF-β signaling in the Ctrl, MDA5+, and MDA5- groups. (D) Sankey plot showing the secreting and receiving niche of the TGF-β signaling. (E) Heatmap showing the normalized pathway activity scores calculated by PROGENY in all samples. (F) Normalized activity scores of the JAK-STAT pathway in representative samples(left). Violin plot showing the comparison of JAK-STAT activity scores in the Ctrl, MDA5+, and MDA5- groups (right-top). Violin plot showing the JAK-STAT activity scores in each niche type of MDA5+ muscles (right-bottom). (G-J) The above analyses were performed for the TGF-β(G), NF-κB (H), Hypoxia (I), and TNF-α (J) pathways (*, p-value < 0.05; **, p-value < 0.01; ***, p-value < 0.001).

To strengthen the above results, we examined the downstream gene expression of the signaling pathways by PROGENY analysis (Fig. 5E, Suppl Table S5). As a result, JAK-STAT pathway activities were upregulated in the 3 MDA5+ and the CADM47, DM81 muscles, and the MDA5+ muscles showed significantly higher activity compared to MDA5- muscles (Fig. 5E, F, Suppl Fig. S5B). Moreover, consistent with findings from the COMMOT analysis, the activity of the TGF-β pathway was upregulated in the MDA5+ muscles, together with increased NF-κB, hypoxia, TNF-α, PI3K, VEGF, and p53 pathways compared to MDA5- muscles (Fig. 5G-J, Suppl Fig. S5B, C). Furthermore, we examined the pathway activities in each niche type. Notably, the activities of TGF-β, hypoxia, NF-κB, and TNF-α pathways were enriched in the EC niches of MDA5+ muscles compared to the other myonuclei niche types (Fig. 5G-J), which is consistent with the result showing increased immune responses and dampened oxygen/gas transport function in the EC niches (Fig. 4E). Taken together, the above findings revealed the unique enrichment of TGF-β, NF-κB, hypoxia, and TNF-α signaling pathways in the EC niches of MDA5+ muscles.

### Macrophages promote humoral response in MDA5+ muscles

The infiltration of immune cells in myositis muscles has been well documented^48–52^, but it is still unclear how the cells promote inflammation and muscle atrophy/damage. To dissect the functional role of immune cells within specific muscle niches, we ranked the cell type abundance of all the Visium spots and defined the “immune cell spots” according to the highest immune cell type abundance (5%). For example, spots with the highest 5% of MΦ1 abundance were classified as “MΦ1 spots” (Fig. 6A). We then quantified the proportion of these spots in each sample and characterized their associated niche types (Fig. 6B). A higher proportion of MΦ1 spots was found in the 2 MDA5+ (CADM42, CADM76) and DM81 muscles. When comparing MDA5+ vs MDA5- muscles, the percentage of MΦ1 spots increased by 2-fold (Fig. 6C), and this increase mainly stemmed from the sMN niches. This expansion of immune infiltrates in MDA5+ muscles was not restricted to macrophages; we also observed higher percentages of spots of plasma cells, B cells, Treg cells, and LAMP3+ DC cells (Suppl Fig. S7A). The CADM subtype similarly showed elevated proportions of these immune cell spots compared to other myositis muscles (Fig. 6D, Suppl Fig. S7B).

**Figure 6.**
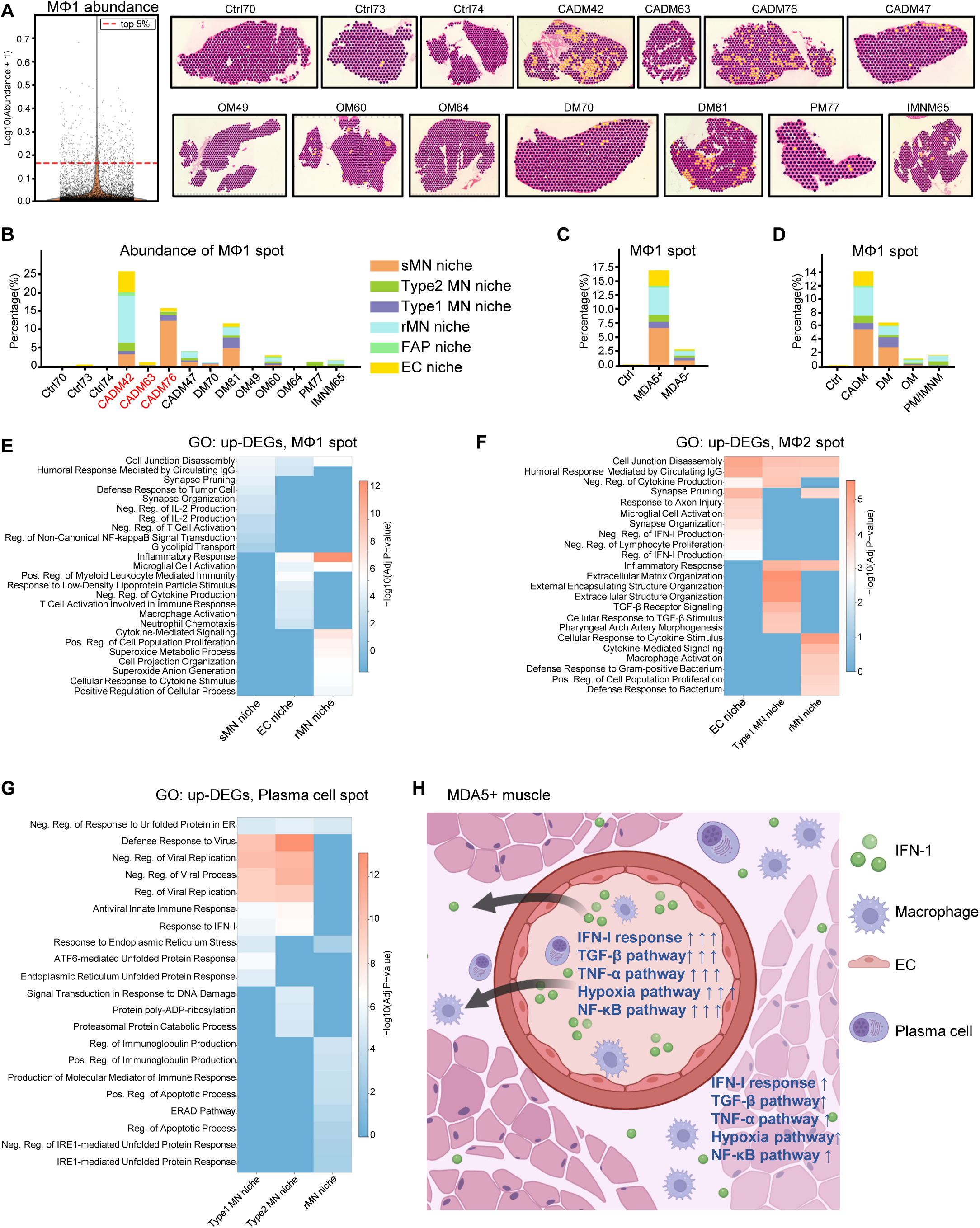
Macrophages promote humoral response in MDA5+ muscles. (A) Left: Violin plot showing the distribution of MΦ1 abundance across all Visium spots, with data pooled from all samples. The top 5% were defined as “MΦ1 spot”. Right: The distribution of the MΦ1 spot in each sample. (B) Stacked bar plot showing the proportion of MΦ1 spots within each sample, stratified by the niche type of the spots. Each bar represents one sample. Bar segments (colors) indicate the proportion of MΦ1 spots belonging to each niche type. (C) Comparison of the proportions of the MΦ1 spots in the Ctrl, MDA5+, and MDA5- groups. (D) Comparison of the proportions of MΦ1 spots in each myositis subtype. (E) Heatmap showing the top 10 enriched GO terms of upregulated genes of MΦ1 spot in the EC niche, rMN niche, and Type2 MN niche, colored by–log10 P value. (F) Heatmap showing the top 10 enriched GO terms of upregulated genes of MΦ2 spot in the EC niche and Type 1 MN niche, colored by–log10 P value. (G) Heatmap showing the top 10 enriched GO terms of upregulated genes of Plasma cell spot in the Type 1 and Type 2 MN niche, colored by–log10 P value. (H) Schematic summarizing the study. Response to IFN-I is upregulated in the EC niches of MDA5+ muscles, possibly activated by the systemic IFN-I from blood. Pathway activities of TNF-α, NF–κB, hypoxia, and TGF–β are also enriched in the EC niches. Immune cells, such as macrophages and plasma cells, infiltrate into muscles via blood vessels.

To elucidate the functional impact of these immune cells, we compared gene expression of the immune cell spots vs. non-immune cell spots in each niche type of MDA5+ muscles. For example, we compared MΦ1 spots vs non-MΦ1 spots in the sMN niche of MDA5+ muscles, and found upregulated pathways, including “humoral immune response mediated by circulating IgG” and “regulation of non-canonical NF-κB signal transduction” enriched in the MΦ1 spots. Similarly, MΦ1 spots in the EC niches and rMN niches showed upregulated pathways of “humoral immune response mediated by circulating IgG”, “cytokine-mediated Signaling”, “T cell activation involved in immune response”, and “inflammatory response”, suggesting that MΦ1 may promote the humoral response to autoantibodies and immune cell activation in MDA5+ muscles (Fig. 6E). The MΦ2 spots in the EC niche, Type 1 MN niche, and rMN niche also showed upregulated pathways of “humoral immune response mediated by circulating IgG” (Fig. 6F), indicating a broader role for macrophages in mediating humoral immunity. Furthermore, antiviral response, immunoglobulin production, and endoplasmic reticulum stress-related pathways were found to be upregulated in the plasma cell spots (Fig. 6G). Moreover, we found Treg cell spots in the Type 1 MN niche associated with negative regulation of immune response, and LAMP3+ DC cell spots in the Type 2 MN niche were associated with upregulated IFN-I and antiviral response (Suppl Fig. S7C, D). Collectively, the above findings illuminate the functional impact of spatially localized immune cells, highlighting a unique pathogenic mechanism in MDA5+ muscles wherein macrophages promote humoral immune response.

## Discussion

MDA5+ myositis is a distinct subtype of myositis characterized by scarce or absent muscle involvement and prominent involvement of skin and lungs. To enhance our understanding of the molecular underpinnings of MDA5+ myositis, we profiled the spatial transcriptome of muscle biopsies collected from a small cohort of myositis patients composed of various subtypes and serotypes. The UMAP clustering separated the MDA5+ muscles well from the healthy muscles, suggesting a distinct spatial transcriptomic profile of the MDA5+ muscles characterized by an IFN-I signature (Fig. 1). Next, we analyzed the cell type composition and found an unexpected increase in immune cells and decreased stressed MN in the MDA5+ muscles (Fig. 2). This increased immune cell infiltration could be explained by immune responses and damage to blood vessels (Fig. 4), leading to local inflammation and enhanced infiltration of immune cells through the wall of the vessel into the perivascular regions. In line with this, we found that staining for CD44, a marker for immune cells that also functions in cell migration as an adhesion molecule, was also enriched at the vascular regions. Unlike IBM or IMNM, in which the CD8+ cytotoxic cells directly target muscle fibers, the increased immune cells are focal in the vascular region of MDA5+ muscles, which may not directly target muscle fibers, explaining why fewer stressed MN were detected in MDA5+ muscles. Furthermore, we found the stressed MN colocalized with aerobic Type 1 fibers, which are more sensitive to hypoxia, possibly caused by EC damage, compared to anaerobic Type 2 fibers.

Spatial transcriptomics overcomes the limitations of bulk sequencing by resolving gene expression within specific tissue regions/niches. In MDA5+ muscles, our data provides the first transcriptome-wide view of six characteristic niches, with a particular focus on the EC niche (blood vessel region). Within this region, we discovered a cohort of IFN-I-induced genes that were upregulated compared to the myofiber region (Fig. 4). Concurrently, we identified a downregulation of oxygen transport-related genes, pointing to significant vascular dysfunction. More interestingly, we found barely detectable IFN mRNA expression in myositis muscles, suggesting that the IFN may originate from the blood; the blood vessels respond first to the circulating interferon, before muscle fibers respond to the interferon diffused from the blood vessels. Consistent with this hypothesis, we found that the expression of the identified group of IFN-I-stimulated genes was upregulated in the EC niches to a greater degree than in the myonuclei niches. Interestingly, ECs in the skin biopsies of DM patients showed strong IFN-I inducible MX1 and ISG15 staining^8, 9, 53^, suggesting this EC response to IFN-I is a systemic phenomenon.

Moreover, we analyzed the cell-cell communication and pathway activity and found that hypoxia, NF-κB, TNF-α, and TGF-β pathway activities showed enrichment in the EC niche of the MDA5+ muscles. Consistent with our findings, positive staining of the phosphorylated p65 subunit of NF-κB was found in the blood vessel endothelium of DM patients^54^. Immunoreactive TNF-α granules were also detected on ECs at perifascicular sites in DM muscles^55^. TGF-β has been reported to be upregulated in the serum of DM/PM patients complicated with ILD^56^, as well as in the muscle biopsies of DM and IBM patients^57, 58^. Consistently, our data revealed the enriched response to TGF-β signaling in the EC niche in MDA5+ muscles (Fig. 5G). Interestingly, a mouse study constitutively activating TGF-β in the ECs resulted in severe cutaneous, pulmonary, and microvascular fibrosis, and loss of ECs due to EndoMT, mimicking the manifestations of MDA5+ DM^59^, suggesting the role of the TGF-β pathway in promoting MDA5+ myositis.

Finally, we analyzed the important role of immune cells in myositis pathogenesis. In the MDA5+ muscles, macrophage spots were associated with upregulation of humoral response to circulating IgG, immune cell activation/proliferation, suggesting that macrophages may play an important role in responding to autoantibodies and promoting inflammation in MDA5+ patients. This is consistent with previous reports showing that macrophage activation correlates with disease activity and poor prognosis of MDA5+ patients^6^.

In summary, our results reveal unique features of MDA5+ muscle and identify several potential therapeutic targets for MDA5+ myositis (Fig. 6H). These include the upregulated IFN-I–induced gene program, NF–κB, TNF–α, and TGF–β pathways in the pericyte–EC niche, as well as infiltrating macrophages and plasma cells. Supporting this, therapies targeting related pathways have shown promise: JAK inhibition (tofacitinib) in refractory MDA5+ myositis^6^; anti–TGF–β (pirfenidone) in MDA5–associated ILD^60^; and B–cell depletion (rituximab or CD19 CAR–T) in MDA5+ RP–ILD^61, 62^. IVIG(Intravenous Immunoglobulin), which neutralizes plasma cell–derived autoantibodies, also improves survival^63^. TNF–α inhibition in other myositis subtypes has yielded mixed outcomes^64^, and NF–κB blockade is being explored in other autoimmune and muscular diseases; future studies should evaluate these strategies specifically in MDA5+ myositis.

We must point out that our study was limited to a cohort of only 11 patients due to the rarity of myositis. The sample size may not be sufficient to investigate the pathogenesis of IMNM or PM subtypes, or to dissect serotypes other than MDA5+. Future studies with larger cohorts are needed to explore these subgroups of myositis. Additionally, the resolution of our spatial transcriptomics data obtained from the 10x Visium_V2 platform was not at the single-cell level, preventing us from delineating gene expression and signaling pathways within individual cells. Nevertheless, building on our findings, several critical avenues for future research emerge. First, the application of single-cell resolution spatial transcriptomics would precisely delineate the cellular changes and signaling pathways within the key vascular and fiber niches we have identified. Second, our inability to detect IFN mRNA within muscle tissues, despite a robust IFN-I signature, suggests an extra-muscular source. It is therefore imperative to determine whether circulating leukocytes (PBMCs) or other affected tissues, such as skin and lung, are the primary contributors of systemic IFN in MDA5+ patients. Furthermore, to obtain a comprehensive understanding of this systemic autoimmune disease, spatial transcriptomic profiling of lung and skin lesions from MDA5+ patients is essential. Such studies would directly reveal the shared and tissue-specific pathways, particularly the systemic alterations in vascular pathology. These mechanistic insights will directly inform therapeutic development.

## Methods

### Patient muscle biopsy collection

The study was reviewed and approved by the Joint Chinese University of Hong Kong-New Territories East Cluster Clinical Research Ethics Committee (CREC Ref. No.: 2022.366). All studies were performed in accordance with standard operating procedure and the principles of the Declaration of Helsinki and ICH Good Clinical Practice. Patients enrolled in this study were admitted to the rheumatology unit of the Prince of Wales Hospital (PWH), New Territories, Hong Kong, China, and fulfilled the 2017 American College of Rheumatology/European League Against Rheumatism (ACR/EULAR) IIM classification criteria of myositis. Informed consent was obtained from all study participants. Conchotome muscle biopsies (4mm punch) were collected by a Pro-Mag Ultra Automatic Biopsy Instrument from patients’ lower limb muscles. Two pieces of biopsies were collected at the same time, one sent for clinical examination and one collected for this study. Patients’ demographic information, clinical examination results, and myositis-specific antibodies (MSAs) and myositis-associated antibodies (MAA) were presented in Suppl Table S1. Healthy control hamstring muscle samples were collected during orthopaedic surgery, and informed consents were obtained in written form from the patients. (Ethical approval granted by the Joint Chinese University of Hong Kong-New Territories East Cluster Clinical Research Ethics Committee (Ref. No.: 2021.255-T).

### Immunofluorescence

Immunofluorescence was performed according to our prior publications^15, 65^. Slides were fixed with 4% PFA for 20 min at room temperature and permeabilized in ice-cold methanol for 10 min at −20 °C. After 4% BSA blocking for 1 hour, the sections were incubated with primary antibodies overnight. The Donkey anti-Mouse IgG (H+L) Highly Cross-Adsorbed Secondary Antibody, Alexa Fluor™ 488(Invitrogen, A-21202), and Donkey anti-Rabbit IgG (H+L) Highly Cross-Adsorbed Secondary Antibody, Alexa Fluor™ 594 (Invitrogen, A-21207) were used as secondary antibodies. Primary antibodies and dilutions were used as follows: CD44(Abcam, ab316123), IFITM3 Rabbit mAb 1:150 (Abclonal, A24337), BST2 Rabbit mAb 1:150 (Abclonal, A23445), SAMHD1 Rabbit mAb (Abclonal, A4607), IFI27 1:100(Abcam, ab171919), α-SMA (Invitrogen, Catalog #14-9760-82). Imaging was performed with the Leica 2500DM microsystem.

### Spatial transcriptomics

Fresh muscle tissues were embedded in Optimal Cutting Temperature (OCT), frozen on dry ice, and stored in −80 °C. Specimens were cryo-sectioned at 10μm and placed on a blank Fisherbrand Superfrost Plus Microscope Slide (samples of two individuals were placed on the same slide within a 6.5mm x 6.5mm area). After fixation with chilled MeOH at −20 °C for 30 min, Hematoxylin & Eosin (H&E) staining was performed, following the Visium CytAssist Spatial Gene Expression for Fresh Frozen – Methanol Fixation, H&E Staining, Imaging & Destaining protocol (10x genomics, CG000614-Rev A). Tissue slides were imaged under a microscope, and the ones with the best quality (no bubbles, even H&E staining, no folded/overlapped tissue) were chosen to proceed with destaining using 0.1N HCl. The library preparation followed the user guide of Visium CytAssist Spatial Gene Expression Reagent Kits (10x genomics, CG000495, Rev E). Briefly, after destaining, the tissue slides were incubated with the Human WT Probes v2 overnight (20 hours) for hybridization. After ligation, the probes were released and transferred using the Visium CytAssist machine, which facilitates the transfer of transcriptomic probes from standard glass slides to the Visium capturing slides (2 6.5mm x 6.5mm capture area). After extension, probes were eluted from the Visium slides, and pre-amplification of 10 PCR cycles was performed. The final cycle number of the library was determined by quantitative PCR. Libraries were sequenced on the Illumina NovaSeq-6000 system platform, and 25GB of data were obtained for each capture area.

### Spatial transcriptomics data processing

After the sequencing was completed, the 10x Genomics Space Ranger (v3.0.0) count function was used to align raw fastq files with the human reference transcriptome (GRCh38-2020-A), tissue detection, fiducial detection and barcode/UMI counting. Subsequent analysis was conducted using Python (v3.9.19). Filtered counts were loaded using scanpy (v1.10.3) ^66^ and spots with fewer than 1000 counts, fewer than 200 features, or more than 20% mitochondrial genes were excluded from downstream analysis. After samples were merged, counts per million (CPM) normalization using built-in function sc.pp.normalize_total() and a log transformation using built-in function sc.pp.log1p() were applied to the merged object. sc.pp.highly_variable_genes() function was then used to select highly variable genes with following parameter settings: min_mean=0.0125, max_mean=3, min_disp=0.5. Following the selection of highly variable genes, the sc.pp.scale() function was used to scale the data to zero mean and unit variance. Next, the merged object was subjected to Principal Component Analysis (PCA) using the sc.tl.pca() function with 50 components, and Harmony (v0.1.8) ^67^ was used to eliminate the batch effects. Finally, the merged object was clustered using the Leiden algorithm at resolution 0.5 and plotted using built-in function pl.umap().

### Cell type deconvolution

Spatial data was deconvoluted using Cell2location (0.1.4) ^17^ with 10 cells per location and a detection α of 20. Unnormalized, single-cell data with cluster information were used as a reference, and q05 abundance counts were used for downstream analysis. Correlations between cell type abundances were calculated using the scipy (v1.13.1)^68^ Pearson’s correlation coefficient. The spatial distribution of multiple cell types was visualized in a unified panel through the built-in plot_spatial() function provided by Cell2location.

### Niche identification

Non-negative-matrix factorization (NMF) function of the Cell2location package was applied to the q05 estimation of cell type abundance to identify cellular niches. We calculated 2 to 30 factors and found 6 factors to best represent cellular niches. Each of the six resulting cellular niches was assigned a descriptive name based on its distinctive cell type composition.

### Differentially expression gene (DEG) and GO analysis

Differentially expression gene (DEG) analysis was applied to the merged spatial object, which was normalized to counts per million (CPM) and log-transformed using function sc.tl.rank_genes_groups() on each specific group. Unless otherwise specified, differentially expressed genes (DEGs) were defined using the following thresholds: absolute log2 fold change (logFC) > 1.3, p-value (pval) < 0.05, and fraction of cells expressing the gene in the target group (pts) > 0.1. An exception was made for the analyses presented in Figure 1 and its associated Suppl Figure 1, where a more stringent logFC cutoff of > 2 was applied. GO enrichment analysis was performed on the up-regulated and down-regulated DEGs in each group using built-in function enrichr() provided by GSEAPY (v 1.1.4) ^69^, with the gene set ‘GO_Biological_Process_2025.gmt’ as the reference ^70^.

### Cell-cell communication analysis by COMMOT

For spatial data, COMMOT (v0.0.3) ^71^ was used to construct the Cell-cell communication network using the built-in function ct.tl.communication_direction(), which can provide us the information of the spatial direction of specific signaling pathways or Ligand-Receptor (LR) pairs. Built-in function ct.tl.communication_deg_detection() and ct.tl.communication_impact() were used to obtain the putative genes that may impacted by specific signaling pathways or Ligand-Receptor pairs.

### Pathway activity inference by PROGENy

Pathway activities were inferred from the spatially resolved transcriptomic data using the PROGENy method, implemented via the decoupler package (v2.1.1) ^72^. The analysis was performed with the function dc.op.progeny(organism=“human”) to generate the pathway model, followed by application of the multivariate linear model dc.mt.ulm() on the preprocessed spatial data. Pathway activity scores for each spot were subsequently extracted using dc.pp.get_obsm (key=“score_ulm”).

### Functional analysis of immune spots

In the spatial data, spots with the highest 5% abundance for a given immune cell type across all samples were defined as high-abundance spots for that type, collectively referred to as ’immune spots.’ For instance, spots in the top 5% for MΦ1 were termed “MΦ1 spots.” The same nomenclature was applied to other cell types (e.g., “T cell spots,” “B cell spots”). Following the “immune spots” annotation, DEG analysis was performed to compare the transcriptomes between immune spots (as defined above) and non-immune spots within each niche type. Genes meeting the thresholds of logFC > 1.3, pvals < 0.05, and pts > 0.1 were identified as DEGs. These DEGs were then divided into upregulated and downregulated sets for subsequent GO) enrichment analysis.

### snRNA-seq data analysis

The snRNA-seq dataset from a published paper was used^73^. Nuclei fastq files were trimmed and aligned to the human GRCh38-2024-A transcriptome using the 10x Genomics Cell Ranger (v8.0.1) count function ^74^. The R package Seurat (v5.0.2) ^75^ was then used to perform the QC, normalization, dimensionality reduction, and clustering on each sample. Nuclei with fewer than 1,000 or more than 20,000 counts, fewer than 800 features, or more than 5% mitochondrial genes were defined as low-quality nuclei and excluded from downstream analysis. Data was normalized using SCTransform ^76^ as implemented by Seurat. Subsequently, visualization and clustering were performed using built-in functions in Seurat, including RunUMAP(), FindNeighbors(), and FindClusters(). Next, low-quality clusters were identified using the built-in function FindAllMarkers(), characterized by the absence of marker genes, and the presence of mitochondrial genes or ribosomal genes, and subsequently removed. After excluding all low-quality nuclei, SoupX (v1.6.2) ^77^ was employed to eliminate background mRNA contamination. Following the exclusion of all low-quality nuclei and background mRNA contamination, DoubletFinder (v2.0.4) ^78^ was used to detect and remove putative doublets on each sample. Subsequently, all samples were integrated, and batch effects were corrected using Seurat CCA algorithm. Finally, normalization, dimensionality reduction and clustering were applied on the integrated object and the built-in function FindAllMarkers() was used to identify the marker genes for each cluster, thus enabling cell-type identification.

### Cell-cell communication analysis of snRNA-seq data

The CellChat2 database (restricted to secreted signaling) and R package ^79^ with default parameters were utilized to separately identify biological interactions among different cell types in the same niche using the built-in function aggregateNet(). In this step, all cell types in the same niche were considered as both sources (ligands) and targets (receptors).

## Data availability

The data that support this study are available from the corresponding authors upon reasonable request. The raw sequence and processed spatial transcriptomics data reported in this paper will be deposited in and publicly available through the Gene Expression Omnibus (GEO) database upon publication of this study.

## Code availability

Custom scripts described in the Methods will be made available upon request.

## Author Contributions

Yulong Qiao performed all the experiments; Gexin Liu analyzed all the high-throughput sequencing data; Michael Tim-Yun Ong assisted in the procurement of the healthy human muscle specimens; Hao Sun supervised computational analyses; Ho So secured the myositis muscle biopsies and associated clinical data. Huating Wang supervised experiments and data analyses; Huating Wang, Ho So, and Yulong Qiao conceived the project. Yulong Qiao, Gexin Liu, Ho So, and Huating Wang wrote the manuscript, with inputs from all authors.

## Acknowledgements

This work was supported by the National Key R&D Program of China to H.W. (project code: 2022YFA0806003); Non-Communicable Chronic Disease-National Science and Technology Major Project of China to H. W. (project code: 2024ZD0530400); The InnoHK initiative of the Innovation and Technology Commission of the Hong Kong Special Administrative Region Government to H.W.; Health and Medical Research Fund (HMRF) from Health Bureau of HK to H.W. (project codes: 10210906 and 08190626); Theme-based Research Scheme (TRS) from RGC to H.W. (project code: T13-602/21-N); General Research Fund (GRF) from Research Grants Council (RGC) of the Hong Kong Special Administrative Region, China, to H.W. (project codes: 14108225, 14106521, 14105123, 14103522, and 14105823 to H.W.); the National Natural Science Foundation of China (NSFC) to H.W. (project codes: 82172436); and Area of Excellence Scheme (AoE) from RGC to H.W. (project code: AoE/M-402/20). The Chinese University of Hong Kong (CUHK) Strategic Seed Funding for Collaborative Research Scheme (SSFCRS) to H.W.

## Inventory of Supplementary Information

### 1. Supplementary Figures

**Suppl Fig. S1.**
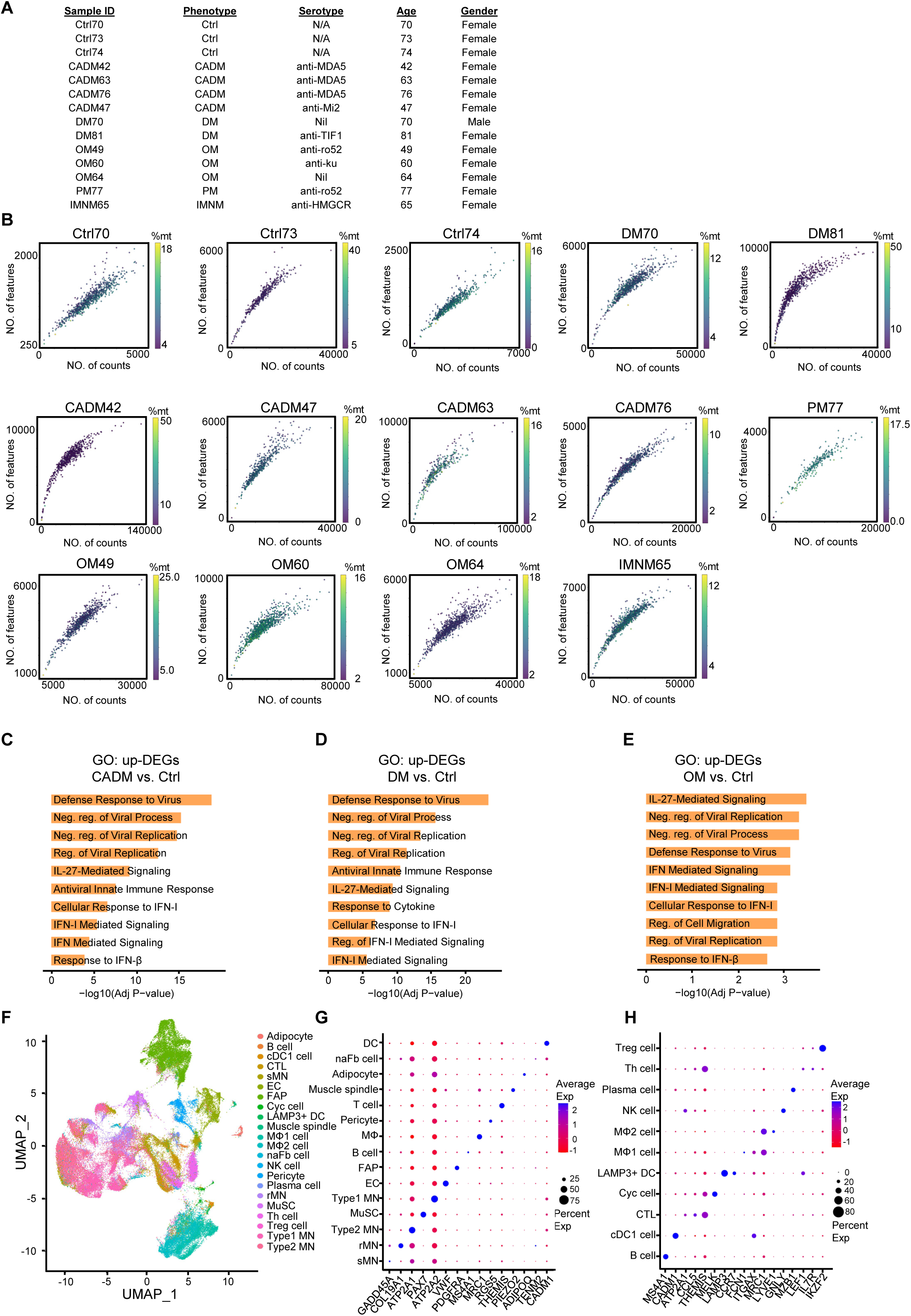
**Clinical information and data quality of spatial transcriptomics. (**A) List of clinical information for each patient. (B) Scatter plots showing quality control (QC) metrics: nFeature, nCount, and percentages of mitochondrial encoded genes (%mt) of each sample. (C-E) GO plot of biological process (BP) terms enriched in CADM group (C), DM (D), and OM (E) groups using the top 200 upregulated genes, compared to the Ctrl group. (F) UMAP plot showing the 22 cell types identified in the reference snRNA-seq data. (G, H) Dot plot showing marker genes of identified cell types.

**Suppl Fig. S2.**
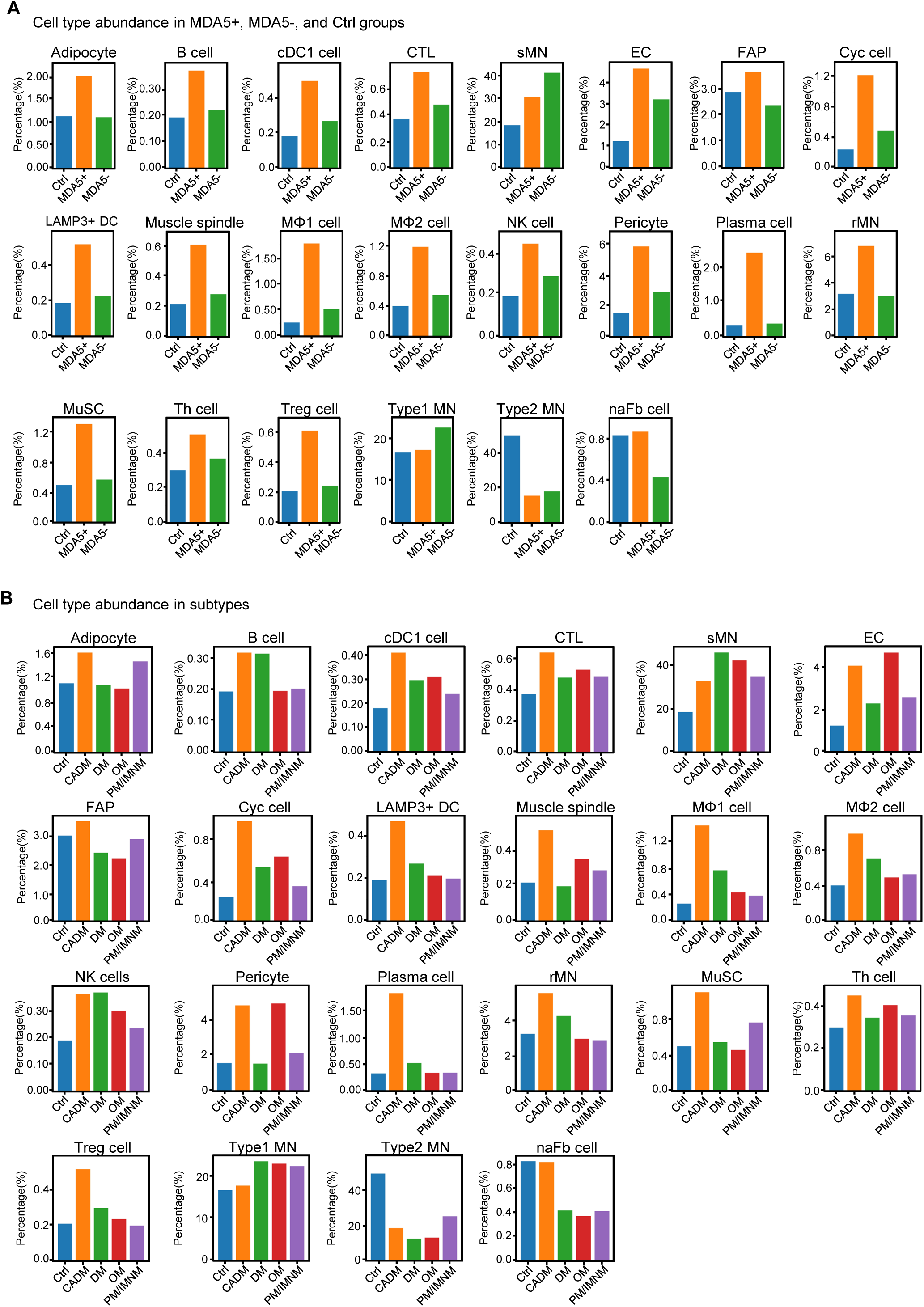
**Quantification of cell type abundance. (**A) Bar plot showing the percentage of each cell type in Ctrl, MDA5+, and MDA5- groups. (B) Bar plot showing the percentage of each cell type in each myositis subtype.

**Suppl Fig. S3.**
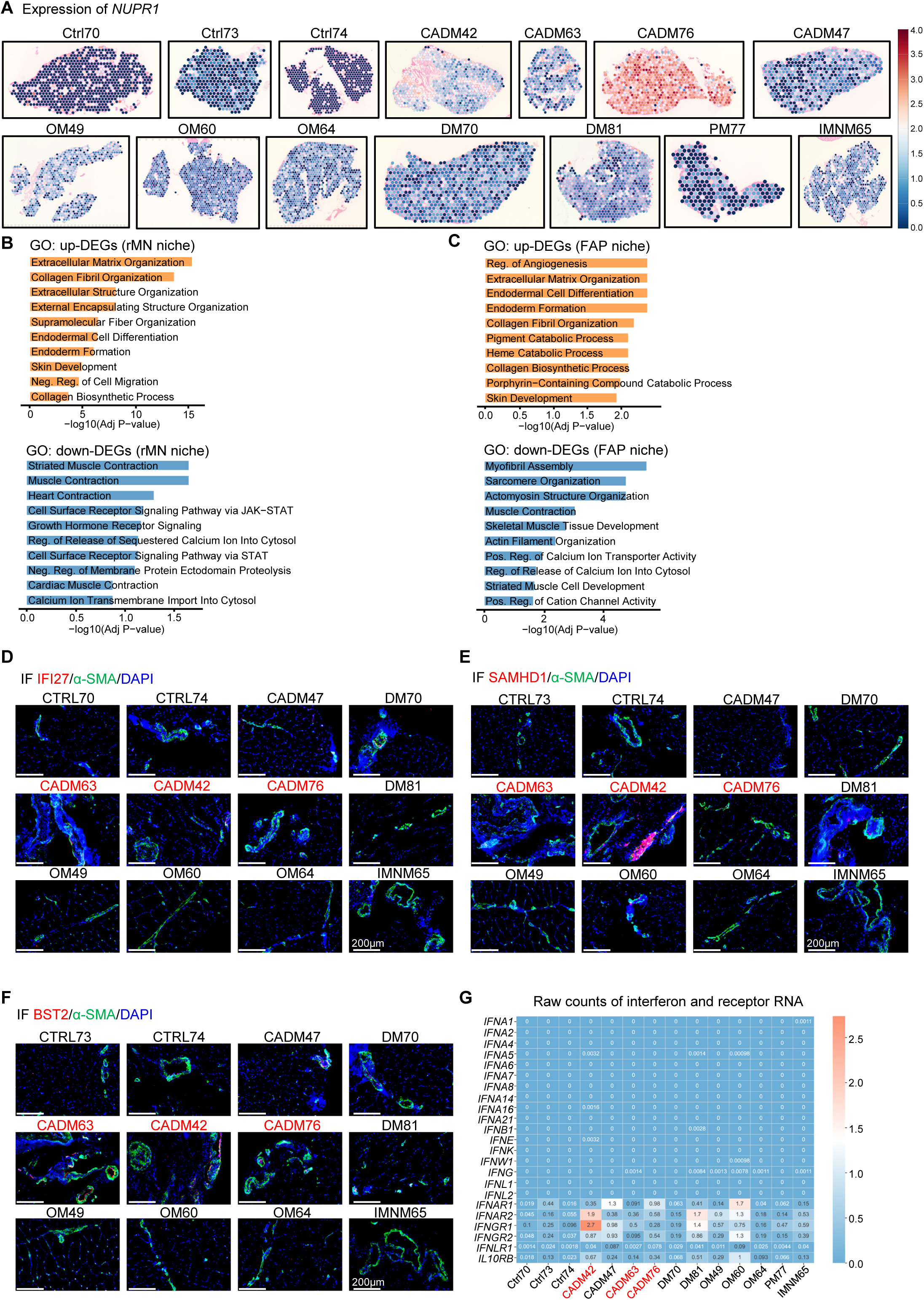
**DEG and GO term enrichment in MDA5+ muscle niches. (**A) Normalized spatial expression of *NUPR1* in all samples. (B-C) Top 10 enriched GO terms for up- and down-regulated genes in the rMN niche (B) and FAP niche (C). (D-F) Immunofluorescence staining of IFI27 (D), SAMHD1 (E), and BST2(F) in representative myositis muscles. Nuclei were stained with DAPI (blue). Blood vessels are labeled with α-SMA (green). Scale bar = 200 μm. (G)Heatmap showing the normalized raw counts of the expression of interferon and interferon receptors in each sample.

**Suppl Fig. S4.**
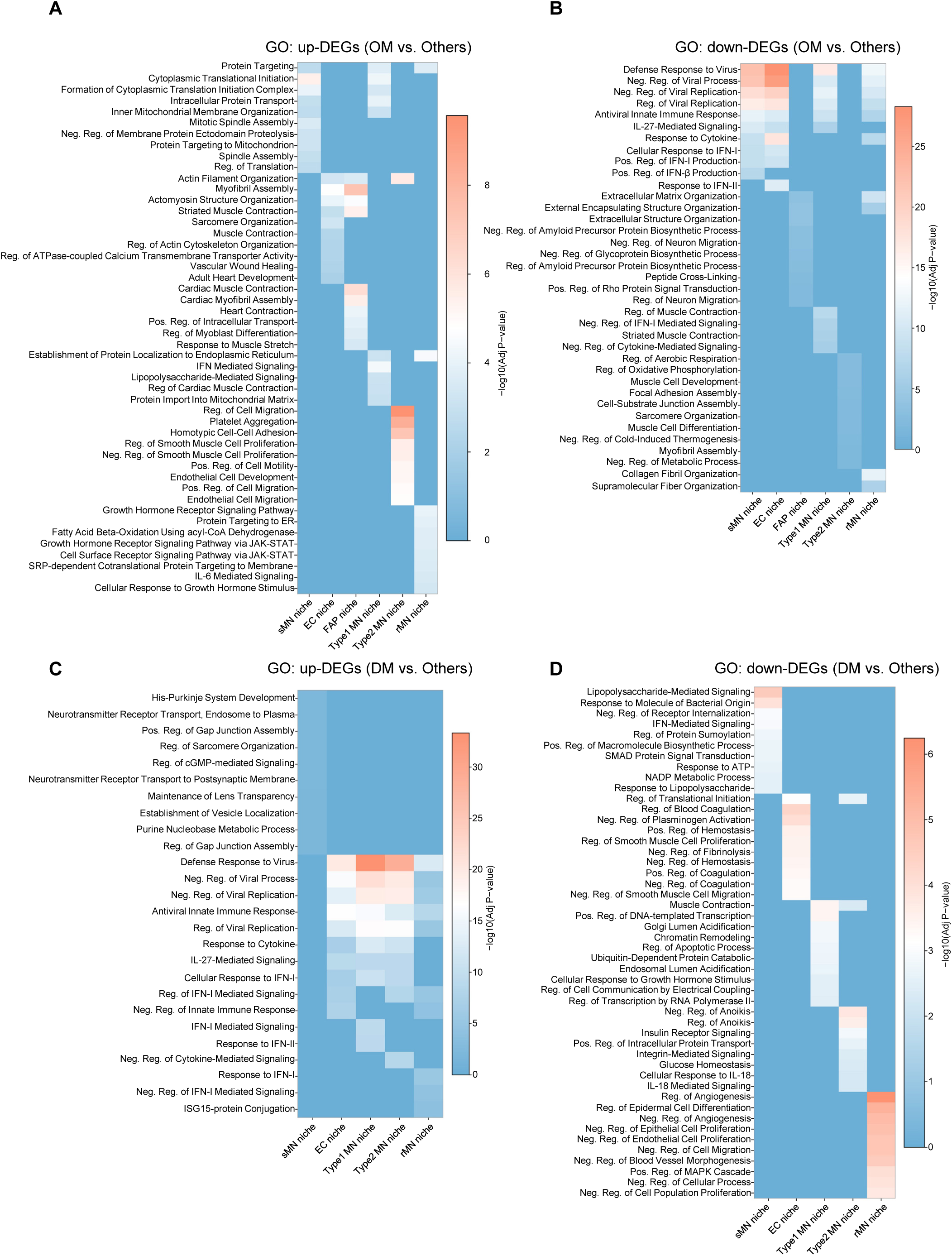
**GO analysis of DEGs in OM and DM subtypes. (**A-B) Heatmap showing the top 10 enriched GO terms of upregulated genes (A) and down-regulated genes(B) in the six niche types of OM, compared to all the other myositis subtypes, colored by–log10 P value. (C-D) Heatmap showing the top 10 enriched GO terms of upregulated genes (C) and down-regulated genes(D) in the six niche types of DM (excluding the FAP niche with no GO terms enriched), compared to all the other myositis subtypes, colored by–log10 P value.

**Suppl Fig. S5.**
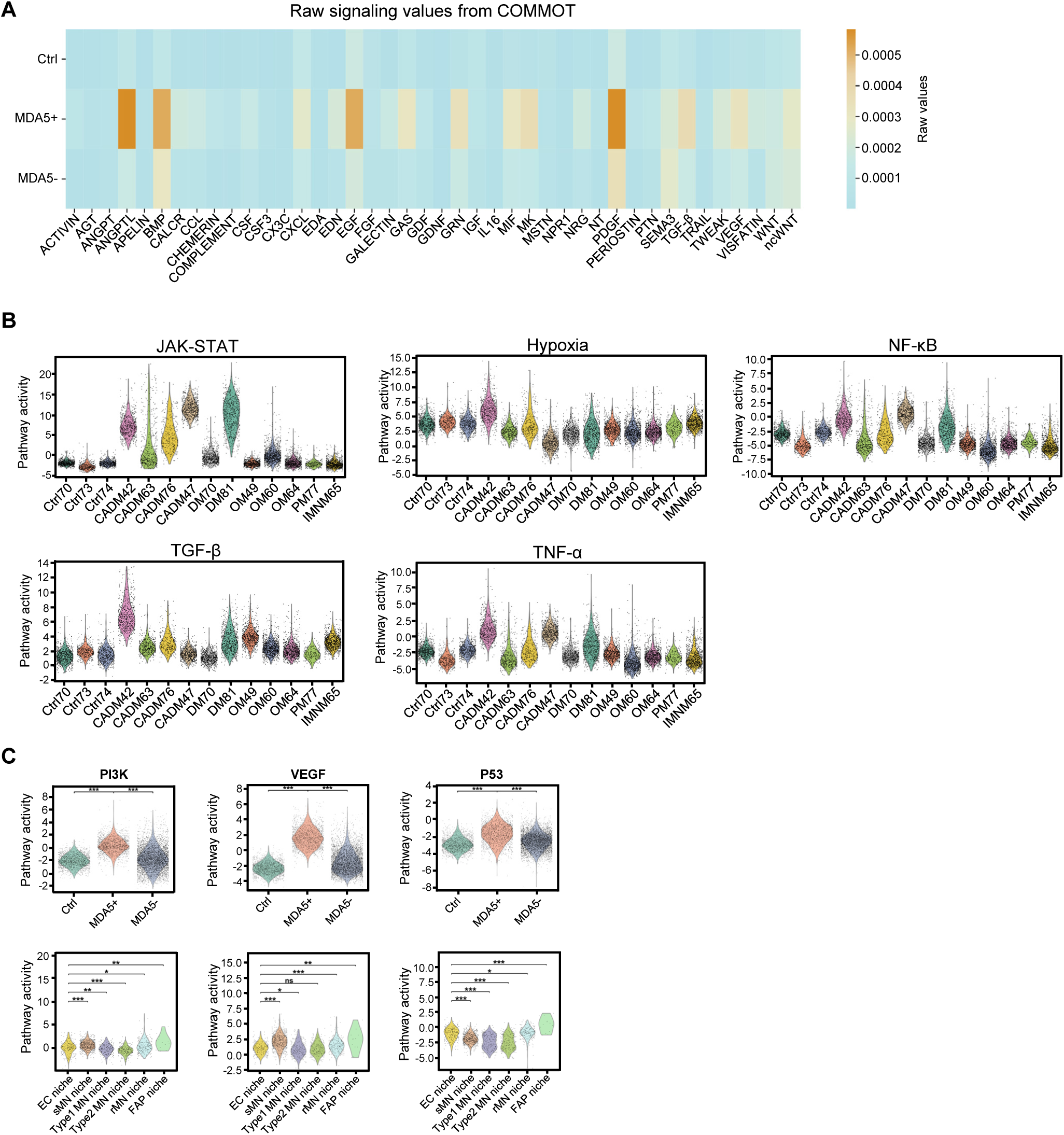
**Signal communication and pathway activities in myositis muscles. (**A) Heatmap showing the normalized signaling communication strength calculated by COMMOT in Ctrl, MDA5+, and MDA5- groups. (B) Violin plots showing the pathway activity of JAK-STAT, Hypoxia, NF-κB, TGF-β, and TNF-α in each sample. (C) Top: Violin plots showing the pathway activity of PI3K, VEGF, and p53 in the Ctrl, MDA5+, and MDA5- groups. Bottom: Violin plots showing the pathway activity of PI3K, VEGF, and p53 in the six niches of MDA5+ muscles.

**Suppl Fig. S6.**
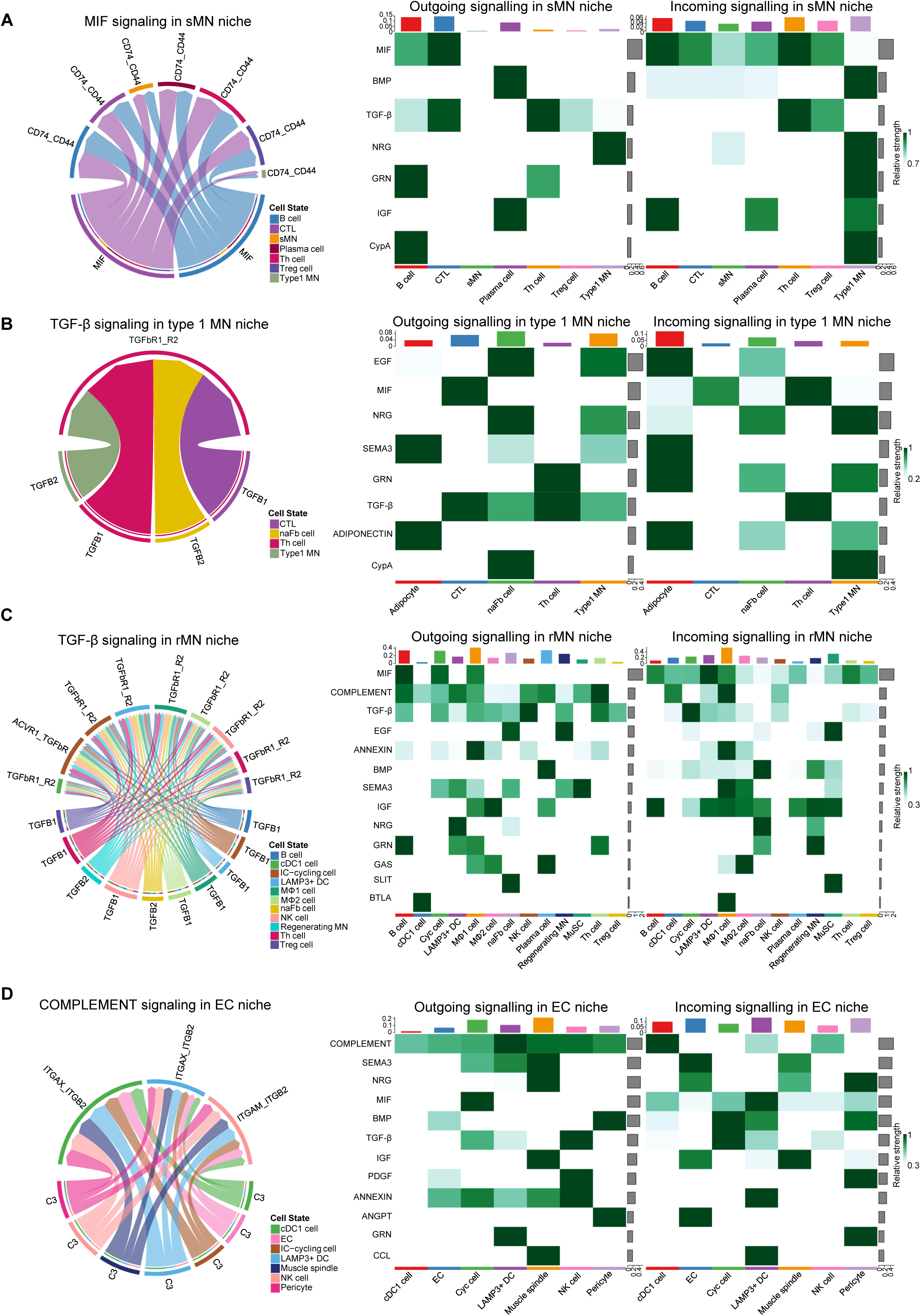
**Cell-cell crosstalk identified from snRNA-seq data. (**A-D) Left: Chord diagram visualizing cell-cell communication through MIF signaling in the sMN niche (A), TGF-β signaling in the Type 1 MN niche (B), TGF-β signaling in the rMN niche (C), and COMPLEMENT signaling in the EC niche (D). Right: Heatmaps showing the strength of all the outgoing and incoming signaling between the cell types in the sMN niche(A), Type 1 MN niche(B), rMN niche(C), and EC niche(D).

**Suppl Fig. S7.**
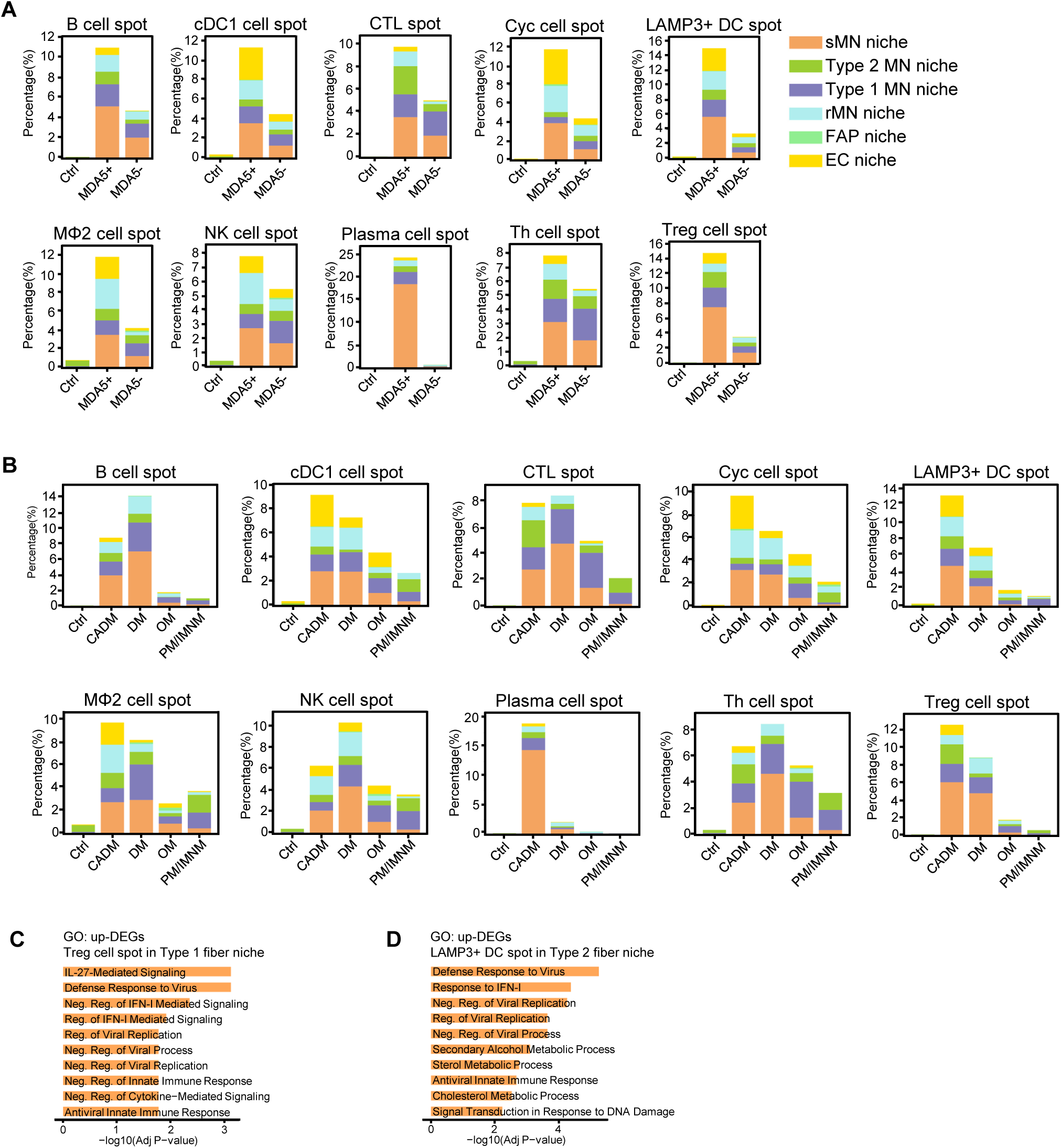
**Quantification and analysis of immune cell spots. (**A) Bar plots showing the percentage of the 10 types of “immune cell spots” in the Ctrl, MDA5+, and MDA5- groups. (B) Bar plots showing the percentage of the 10 types of “immune cell spots” in each myositis subtype. (C) Top 10 enriched GO terms for the upregulated genes of the Treg cell spots in the type 1 MN niche, compared to non-Treg cell spots. (D)Top 10 enriched GO terms for the upregulated genes of the LAMP3+ DC cell spots in type 2 MN niche, compared to non-LAMP3+ DC cell spots.

### 2. Supplementary Tables

Suppl Table S1. Clinical information, data quality, DEG, and GO analysis of spatial transcriptomics.

Suppl Table S2. Quantification of cell type proportions.

Suppl Table S3. DEGs of six niche types and proportion of niches across all samples.

Suppl Table S4. Analysis of DEGs and GO terms among subtypes and serotypes for each niche.

Suppl Table S5. Analysis of cellular crosstalk and pathway activities.

Suppl Table S6. DEG and GO analysis of immune cell spots in different niches of MDA5+ muscles.

